# A spatiotemporal ventricular myocyte model incorporating mitochondrial calcium cycling

**DOI:** 10.1101/604926

**Authors:** Z. Song, L-H Xie, J. N. Weiss, Z. Qu

## Abstract

Intracellular calcium (Ca^2+^) cycling dynamics in cardiac myocytes are spatiotemporally generated by stochastic events arising from a spatially distributed network of coupled Ca^2+^ release units (CRUs) that interact with an intertwined mitochondrial network. In this study, we developed a spatiotemporal ventricular myocyte model that integrates mitochondria-related Ca^2+^ cycling components into our previously developed ventricular myocyte model consisting of a 3-dimensional CRU network. Mathematical formulations of mitochondrial membrane potential, mitochondrial Ca^2+^ cycling, mitochondrial permeability transition pore (MPTP) stochastic opening and closing, intracellular reactive oxygen species (ROS) signaling, and oxidized Ca^2+^/calmodulin-dependent protein kinase II (CaMKII) signaling were incorporated into the model. We then used the model to simulate the effects of mitochondrial depolarization on mitochondrial Ca^2+^ cycling, Ca^2+^ spark frequency and amplitude, which agree well with experimental data. We also simulated the effects of the strength of mitochondrial Ca^2+^ uniporters and their spatial localization on intracellular Ca^2+^ cycling properties, which substantially affected diastolic and systolic Ca^2+^ levels in the mitochondria but exhibited only a small effect on sarcoplasmic reticulum and cytosolic Ca^2+^ levels under normal conditions. We show that mitochondrial depolarization can cause Ca^2+^ waves and Ca^2+^ alternans, which agrees with previous experimental observations. We propose that this new spatiotemporal ventricular myocyte model, incorporating properties of mitochondrial Ca^2+^ cycling and ROS-dependent signaling, will be useful for investigating the effects of mitochondria on intracellular Ca^2+^ cycling and action potential dynamics in ventricular myocytes.

## Introduction

Spatiotemporal intracellular calcium (Ca^2+^) cycling dynamics in cardiac myocytes are generated by stochastic phenomena arising from a three-dimensional Ca^2+^ release unit (CRU) network composed of ryanodine receptor (RyR) clusters in the sarcoplasmic reticulum (SR) membrane. The SR is the major Ca^2+^ store in cardiac myocytes, and the CRUs are coupled via Ca^2+^ diffusion in both the cytosolic and SR space. The intracellular Ca^2+^ dynamics that arise from this network include Ca^2+^ sparks, spark clusters, partial and persistent Ca^2+^ waves, and whole-cell Ca^2+^ oscillations, which play important roles in excitation-contraction, signal transduction, and arrhythmogenesis. In addition to the SR, mitochondria also store Ca^2+^ by taking up Ca^2+^ via mitochondrial Ca^2+^ uniporter (MCU) and extruding Ca^2+^ via mitochondrial Na^+^-Ca^2+^ exchange (NCX_m_) or mitochondrial permeability transition pores (MPTPs). It has also been shown that mitochondrial depolarization and other metabolic stress affect the Ca^2+^ cycling dynamics, including Ca^2+^ spark (1, 2), Ca^2+^ alternans (3-8), and spontaneous Ca^2+^ release and waves (9, 10). Mitochondrial depolarization and Ca^2+^ cycling can also affect the action potential dynamics, such as promoting early afterdepolarizations (EADs) (11, 12), and lower membrane excitability by opening of K_ATP_ channels (13-15) to promote arrhythmias, or modulating the pacemaking activity of cardiomyocytes (16, 17).

Mitochondria are coupled to intracellular Ca^2+^ cycling and the cardiac action potential in several ways. First, mitochondria directly influence intracellular Ca^2+^ cycling via mitochondrial Ca^2+^ cycling in which cytosolic Ca^2+^ is sequestered in mitochondria via MCU and then is released back to the cytosol via NCX_m_ or opening of mitochondrial permeability transition pores. Second, mitochondria related reactive oxygen species (ROS) signaling and redox regulation affect RyR open probability and SERCA pump activity (18-23) as well as ion channel open probability. Third, the SERCA pump and other sarcolemmal ion pumps require ATP. Low ATP may impair these pumps and trigger the opening of the ATP-dependent K^+^ channels. Mitochondrial ROS production and SR Ca^2+^ release form a positive feedback loop since leaky RyRs may result in more mitochondrial Ca^2+^ uptake which can cause mitochondrial depolarization and ROS production (24, 25). Moreover, a ventricular myocyte contains thousands of mitochondria, forming a mitochondrial network exhibiting spatiotemporal dynamics (26-33). This network, coupled with the CRU network, also influences spatiotemporal Ca^2+^ cycling dynamics. In addition, the elementary Ca^2+^ release events (e.g. Ca^2+^ sparks) occur randomly due to random L-type Ca^2+^ channel (LCC) and RyR openings (34, 35). Similarly, the mitochondrial membrane potential flickering (36, 37) and ROS flashes (38, 39) also occur randomly in the single mitochondrion level. Therefore, to understand the interactions between mitochondria and intracellular Ca^2+^ cycling and action potential, both the spatial distribution of subcellular components and random behaviors arising from ion channel stochasticity need to be considered.

In this study, we developed a ventricular myocyte model that includes a detailed spatial network of CRUs and mitochondria, which couples to membrane voltage, intracellular cytosolic, SR, and mitochondrial Ca^2+^ cycling. The model also incorporates intracellular ROS signaling, oxidized Ca^2+^/calmodulin-dependent protein kinase II (CaMKII) signaling, stochastic MPTP opening, and mitochondrial Ca^2+^ buffering. Rather than incorporating detailed models of glycolytic and metabolic pathways, we used simplified models for mitochondrial membrane potential, ATP, and ROS described by phenomenological functions. The model was validated against experimental data from literature and predicts Ca^2+^ cycling dynamics such as mitochondrial depolarization induced Ca^2+^ alternans and Ca^2+^ waves. This model provides a new platform to investigate the effects of mitochondria on intracellular Ca^2+^ cycling and action potential dynamics of ventricular myocytes.

## Mathematical modeling and simulation methods

The mathematical equations and parameters of the model were described in detail in *SI*. Here we outline the basics and the major changes we made from previous models.

### The overall ventricular myocyte model structure

The spatiotemporal myocyte model (Fig.1A) consists of a three-dimensional coupled network of CRUs and mitochondria, modified from the model in our recent study (12), which contains 21,504 (64×28×12) CRUs and 5,376 (64×14×6) mitochondria. These numbers can be changed for modeling different cell sizes. The CRU distance is ∼2 μm in the longitudinal direction and ∼1 μm in the transverse directions. According to Vendelin et al. (40), the average distance between mitochondria is ∼2 μm in longitudinal direction and ∼1.5 μm in the transverse direction in cardiac myocytes. Therefore, we attached a mitochondrion to every CRU along the longitudinal direction and every other CRU along the transverse direction, resulting in a 4:1 CRU-to-mitochondrion ratio. RyRs and LCCs were modeled by random Markovian transitions (Fig.1B and C). The major improvements in the model from our previous study (12) are incorporation of: i) a new stochastic MPTP gating model (Fig.1F), a new model for mitochondrial Ca^2+^ buffering, and simplified models for cytosolic ROS and ATP; ii) the MCU formulation by Williams et al. (41) and the oxidized CaMKII signaling formulation by Foteinou et al. (42) (Fig.1D and E); and iii) ATP, ROS and CaMKII signaling effects on RyRs, SERCA pump, and the relevant ionic currents based on previous models and experimental information.

**Figure 1.**
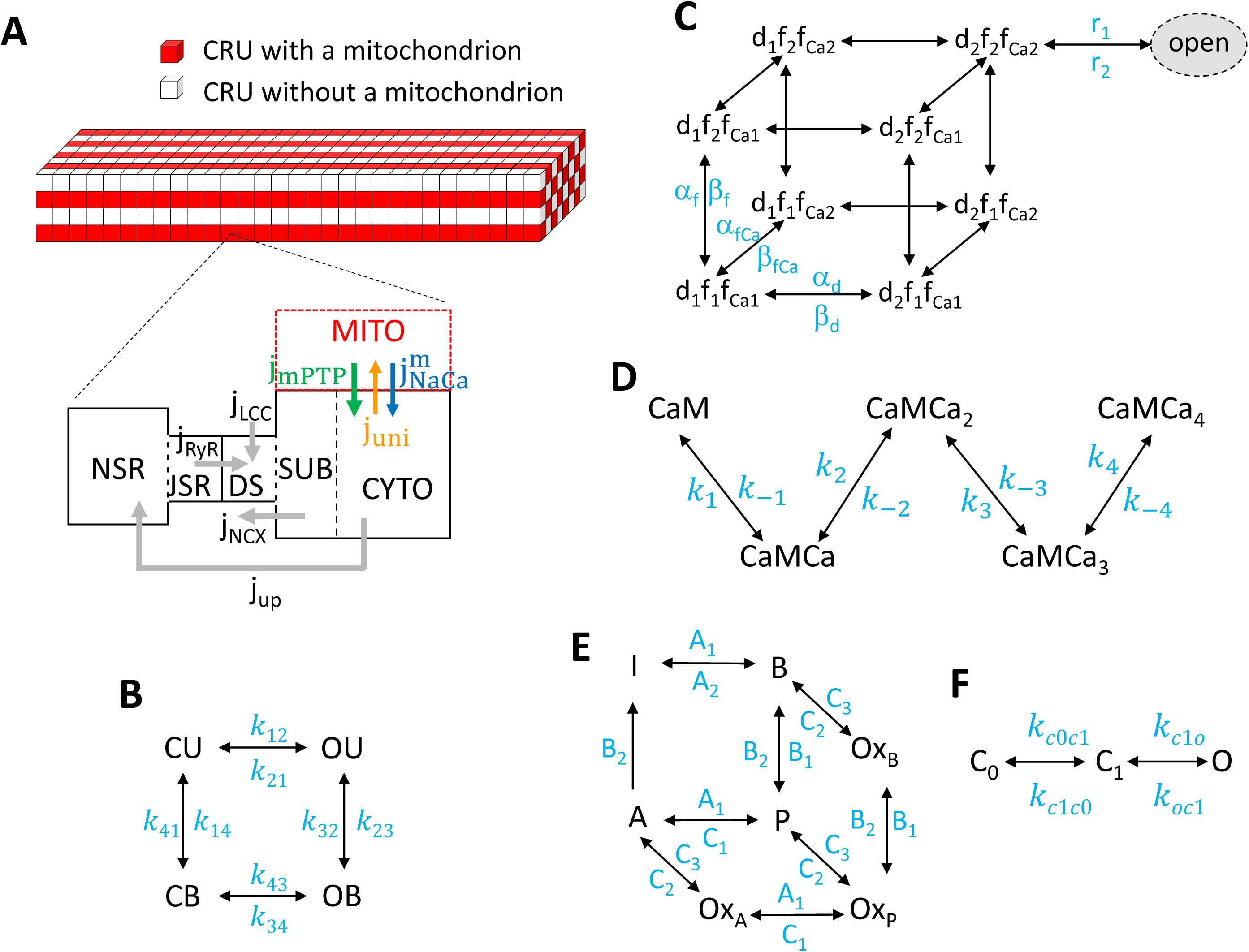
Schematic diagram of the ventricular myocyte model. **A.** Schematic diagram of the CRU-mitochondria network consisting of 64×28×12 CRUs and 64×14×6 mitochondria. Mitochondria are attached to each CRU in the longitudinal direction and every other CRU in the transverse direction. The compartments contained in a CRU are: CYTO—cytosolic space; SUB—submembrane space; DS— dyadic space; JSR—junctional SR; NSR—network SR. Gray arrows indicate the Ca^2+^ fluxes via LCC, NCX, RyRs and SERCA. When a mitochondrion is attached to a CRU, Ca^2+^ fluxes via MCU, NCX_m_ and MPTP are indicated by the colored arrows. **B**. The 4-state RyR model. **C**. The 9-state LCC model. **D**. The 5-state CaM model. **E**. The 7-state CaMKII model. **F**. The 3-state MPTP model.

### Membrane potential of the cell

The membrane potential (V) of the cell is described by the following differential equation:

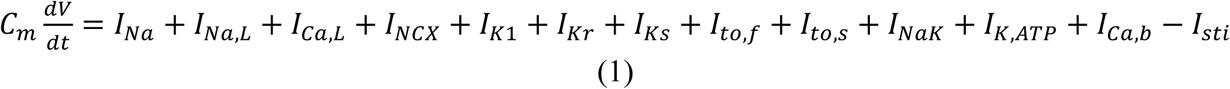

Where *C*_*m*_=1 *μF/cm*^*2*^ is the cell membrane capacitance and *I*_*sti*_=-50 *μA/cm*^*2*^ is the stimulus current density. The ionic current formulations were taken from the rabbit ventricular myocyte model by Mahanjan et al. (43) with modifications, which are summarized below.

#### Na^+^ current

*I*_*Na*_ is the Na^+^ current density and *I*_*Na,L*_ is the late Na^+^ current density, and both of which are CaMKII-dependent using the formulations by Hund et al. (44).

#### L-type Ca^2+^ current

*I*_*Ca,L*_ is the L-type Ca^2+^ current density. Since the stochastic opening of LCCs plays important roles in the stochastic firing of Ca^2+^ sparks, the gating of LCCs was modeled using a 9-state Markovian model (12) to describe the different states of the LCCs (Fig.1C). We added the CaMKII-dependent regulation by slowing the inactivation of the LCCs (45), using the formulation by Hund and Rudy (46).

#### Na^+^/K^+^ pump current

The Na^+^-K^+^ pump current density (*I*_*NaK*_) model was modified from the formulation by Cortassa et al. (47) to incorporate dependencies on ATP and ADP.

#### ATP-sensitive K^+^ current

An ATP-sensitive K^+^ current (*I*_*KATP*_) was added to the model using the formulation by Matsuoka et al. (48).

### Cytosolic and SR Ca^2+^ cycling

We incorporated CaMKII, ATP, and ROS regulation of RyRs and SERCA pumps into the model as described below.

#### RyR open probability

The effects of mitochondrial depolarization on the RyR open probability are mediated by both oxidized CaMKII signaling (49) or redox regulation (20), which increase the RyR open probability. To simulate these effects, we modified the close-to-open rate of RyRs as follows:

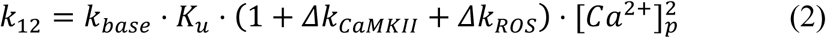

where *Δk*._*CaMKII*_ is the CaMKII-dependent component and *Δk*_*ROS*_ is the ROS-dependent component. *k*_*base*_ and *K*_*u*_ are two rate constants, and [*Ca*^2+^]_*p*_ is the local Ca^2+^ concentration in the corresponding dyadic space.

#### SERCA pump

The formulation by Cortassa et al. (47) was used for the dependence of SERCA pump on ATP. The effects of ROS on SERCA activity are via direct redox regulation (20) which slows the SERCA pump, or via CaMKII phosphorylation of phospholamban which reduces the half-maximum value (*k*_*i*_) to increase SERCA activity. The effects of CaMKII signaling on phospholamban (and thus SERCA activity) were simulated using the formulation by Hund and Rudy (46). Thus, the formulation of SERCA pump is described as follows:

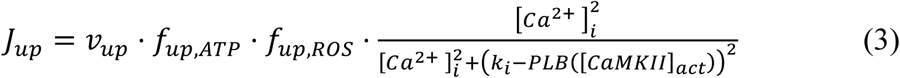

where f_up,ATP_ is an ATP-dependent function taken from Cortassa et al. (47) and f_up,ROS_ is an ROS-dependent function.

#### Background SR leak

We added a CaMKII-dependent background SR Ca^2+^ leak flux following the formulation by Hund et al. (44).

### Mitochondrial membrane potential

Since metabolism is not the focus of this study, we ignored the details for mitochondrial membrane potential (Δψ) generation and usage (50), and used a simple equation to describe Δψ from our previous publication (32), i.e., for each mitochondrion in the cell, Δψ is described by the following differential equation:

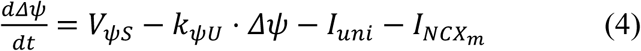

where *V*_*ψs*_ = 3.5 m*V*/*ms* is the Δψ production rate, *k*_*ψU*_ = 0.192 *ms*^−1^ is the usage rate constant, *I*_*uni*_ is the current density via the MCU of the mitochondrion, and 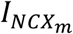 is the current density via its NCX_m_. Note that Eq.4 only holds when the MPTP is closed. MPTP opening is known to effectively dissipate the mitochondrial membrane (51), and therefore, we assume that Δψ is immediately depolarized to zero once the MPTP opens.

A 3-state model of MPTP gating kinetics (Fig.1F) from our previous study (51) was used. The transition rate from the state C_0_ to the state C_1_ was assumed to be Ca^2+^-dependent as follows:

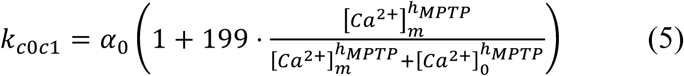

where h_MPTP_ = 5 is the Hill coefficient, [Ca^2+^]_m_ is the free Ca^2+^ concentration in the corresponding mitochondrion, and [Ca^2+^]_0_ is the K_d_ for the half-maximum value. Other transition rates are assumed to be constant.

### Mitochondrial Ca^2+^ cycling

#### Mitochondrial Ca^2+^ uptake

We incorporated the MCU formulation by Williams et al. (41) into the model, which was fitted to experimental MCU fluxes over a wide range of cytosolic Ca^2+^ concentration.

#### Mitochondrial Na^+^-Ca^2+^ exchange

We incorporated the mitochondrial Na^+^-Ca^2+^ exchange formulation by Cortassa et al. (47).

#### Mitochondrial Ca^2+^ buffering

The mitochondrial Ca^2+^ buffering capacity under physiological conditions has been estimated from 33:1 to 30,000:1 (bound/free) (52). Previous studies showed that most matrix Ca^2+^ is buffered by Pi in a pH-sensitive manner, and when MPTP opens, the buffering capacitance is reduced due to the decrease in matrix pH (53, 54). In a recent study (55), the time course of mitochondrial free Ca^2+^ was recorded during MPTP opening induced by NCX_m_ inhibition. After MPTP closure, the mitochondrial free Ca^2+^ re-accumulated with a steep slope, and then exponentially relaxed to a constant slope with a timescale of ∼20 sec. A possible interpretation could be that the buffering capacitance immediately after MPTP closure is still relatively low, and it takes time for the mitochondrial Ca^2+^ buffers to recover to their high capacitance. Therefore, to reproduce this behavior in our model, we assumed that there is a relaxation time constant for mitochondrial Ca^2+^ buffers to adjust their capacitance in response to MPTP closure. Thus, when MPTP opens, the buffering capacitance is set to *β*_*m,on*_, and when MPTP closes, the buffering capacitance relaxes to *β*_*m,off*_ with a time constant τ_m_, i.e.,

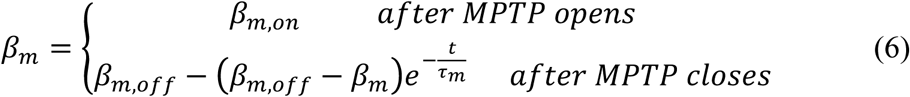

The parameters were chosen to be within the physiological range (52), and were adjusted to reproduce the experimental results by Lu et al. (55) (see Fig.2).

**Figure 2.**
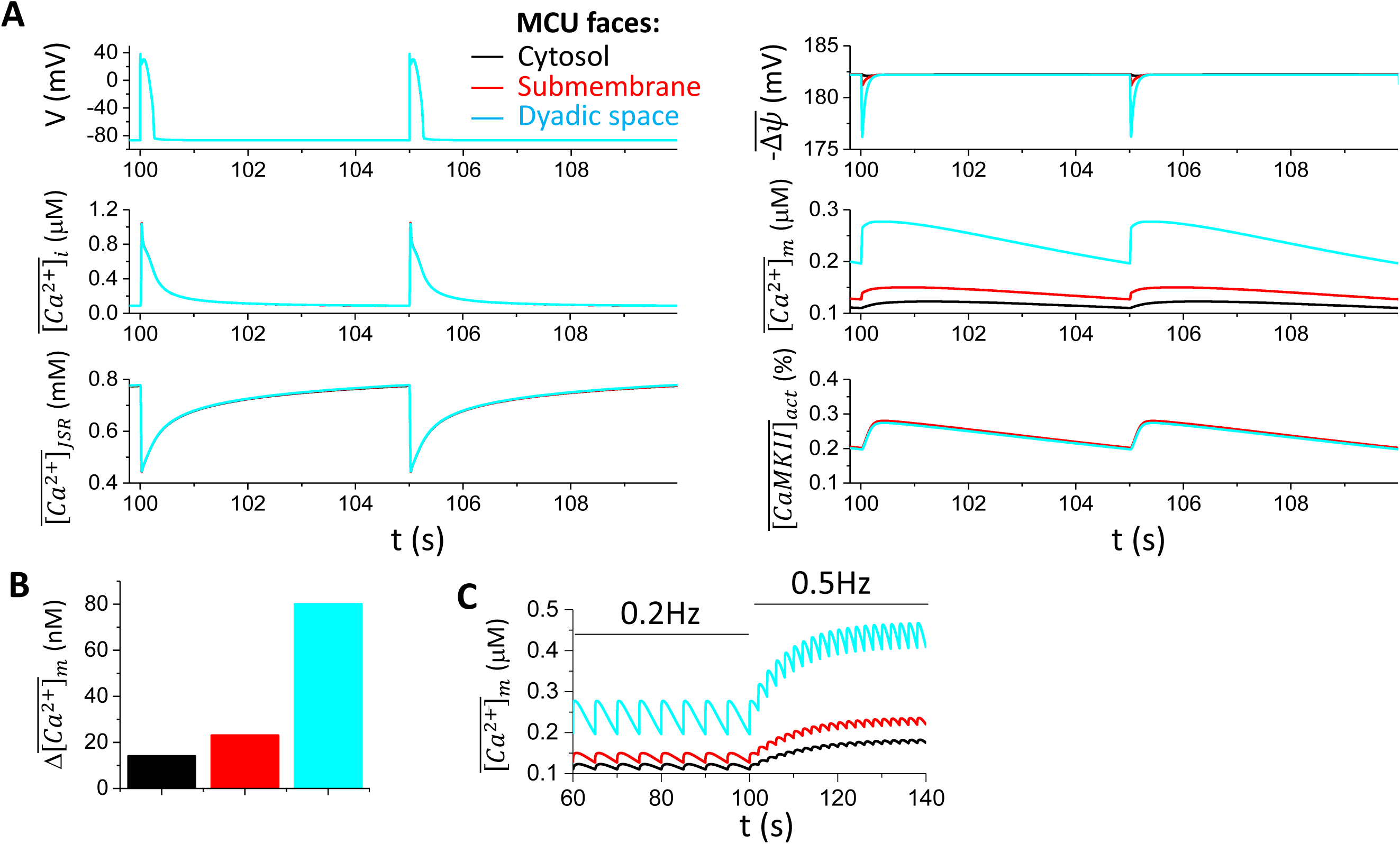
MCU localization on action potential and Ca^2+^ cycling properties under normal conditions. **A.** Time courses of 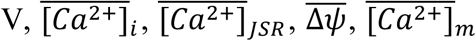, and 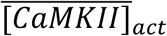 for MCU facing cytosol (black), submembrane space (red), and dyadic space (cyan), respectively. In the model, MCU only senses local Ca^2+^. For example, when the MCU faces cytosol, it only senses the local Ca^2+^ concentration in the corresponding cytosolic space, i.e., [Ca^2+^]_i_. Since MCU localization exhibits small effects on action potential and intracellular Ca^2+^ cycling properties, the colors are overlapped except in the panels for 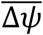 and 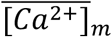. **B**. Mitochondrial free Ca^2+^ amplitude 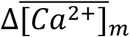 for the three MCU localizations. **C**. Mitochondrial free Ca^2+^ in response to fast pacing. Colored traces correspond to different MCU localizations.

### Cytosolic ROS

Mitochondrial depolarization causes an ROS burst as shown in many experiments (56, 57). In our model, we assume the cytosolic ROS production rate is high when MPTP opens and remains low when MPTP closes. Accurate measurement of cytosolic ROS in cells remains difficult (58), but it is estimated to be less than 0.25 µM under normal conditions and could be 100-fold greater under diseased conditions (59). We applied the following governing equation for the cytosolic ROS dynamics in each CRU:

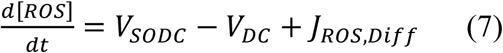

where *V*_*SODC*_ is the mitochondrial ROS production rate, which is a function of mitochondrial membrane potential. If the CRU does not have a mitochondrion attached to it, this term becomes zero. *V*_*DC*_*=K*_*DC*_*[ROS]* is the ROS degradation rate. Based on the fact that the lifetime of H_2_O_2_ is reported to be ∼ 10 ms (60), *K*_*DC*_ was set to be 0.1 *ms*^−1^. *J*_*ROS,Diff*_ is the ROS diffusion flux between neighboring CRUs with the diffusion constant from Yang et al. (32). *V*_*SODC*_ is formulated such that when a mitochondrion depolarizes, the cytosolic ROS level reaches ∼200 *μM* and when mitochondrion remains repolarization, the cytosolic ROS level is ∼0.1 *μM*, which are reasonably within the pathophysiological range (42, 59).

### Cytosolic ATP

ATP cycling in our previous study (50) was formulated using a detailed glycolysis model. Here, since glycolysis dynamics is not the focus of this study, we used a simplified ATP model:

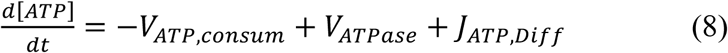

where *V*_*ATPase*_ is the mitochondrial ATP production rate, and *V*_*ATP,consum*_ is the ATP consumption rate. *J*_*ATP,Diff*_ is the ATP diffusion flux between neighboring CRUs with the diffusion constant from Hubley et al. (61). Under the control condition, the cytosolic ATP level is ∼ 4.9 mM.

### CaMKII signaling

The CaMKII signaling model (Fig.1E) developed by Foteinou et al. (42), which includes the oxidation activation pathway, was incorporated into our model.

### Computer simulation

Stochastic transitions of LCCs, RyRs and MPTP were simulated using a modified Gilespie method (62), the differential equations were solved using the Euler method with a time step of 0.01 ms. A time adaptive method (with a time step varying from 0.001 ms to 0.01 ms) was used for computing the action potential upstroke of the ventricular cell and the Ca^2+^ dynamics in the dyadic space. All computer programs were coded in CUDA C with double precision. Simulations were carried out on a high throughput computation cluster consisting of Nvidia Tesla K20c and K80 GPU cards. To simulate an action potential of a PCL 1 sec, it takes ≈ 140 sec of computer time.

## Results

### Effects of MCU localization on whole-cell intracellular Ca^2+^ cycling properties under normal conditions

Mitochondria are tethered in close proximity to SR Ca^2+^ release sites (63), where changes in cytosolic Ca^2+^ concentration are the most dynamic in cardiac cells. For instance, MCUs located close to the dyadic space sense much higher free Ca^2+^ concentration (∼100 *μ*M) than those in the bulk cytosol (∼1 *μ*M). To investigate how the MCU localization affects the intracellular and mitochondrial Ca^2+^ cycling dynamics, we simulated our cell model under normal conditions with MCU facing the bulk cytosol, the submembrane space, or the dyadic space only.

Fig.2A shows the time courses of cell membrane potential (V), whole-cell averaged cytosolic free Ca^2+^ concentration 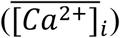, whole-cell averaged free Ca^2+^ concentration in the 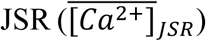, whole-cell averaged mitochondrial membrane potential 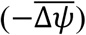, whole-cell averaged free mitochondrial Ca^2+^ concentration 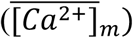, and whole-cell averaged active CaMKII activity 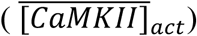 for PCL=5 s and MCU facing different compartments. For all three cases, the action potential, the whole-cell Ca^2+^ transient 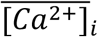, and 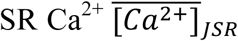, as well as the CaMKII activity exhibit almost no change for MCU sensing Ca^2+^ in different compartments. The mitochondrial membrane potential 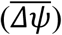 exhibits a small change in which a larger depolarization occurs when MCU senses the dyadic Ca^2+^. Altering MCU localization mainly altered the mitochondrial Ca^2+^ load, with MCU facing the dyad resulting in the highest 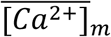 and MCU facing the bulk cytosol the lowest 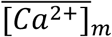. The 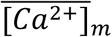 diastolic-to-systolic variations are ∼15%, ∼20%, and ∼40%, respectively, depending on the MCU localization (Fig.2B). This variation is in line with the experiments by Lu et al. (64) who showed that during the steady state pacing (PCL = 5 s), mitochondrial Ca^2+^ concentration 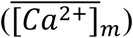 rises to the peak rapidly and declines much more slowly with a 10-20% diastolic-to-systolic variation.

Fast pacing causes the 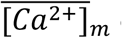 diastolic level to rise with a reduced systolic amplitude (Fig. 2C), replicating the pacing-dependent behavior of mitochondrial free Ca^2+^ reported in experiments by Lu et al. (64). Our simulations predict that MCU localization affects the mitochondrial Ca^2+^ level, which may rise significantly during fast pacing. For instance, if MCU faces the dyadic space, at a PCL of 2 s, the 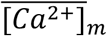 can reach ∼0.4-0.5 µM.

### Effects of MCU activity on the whole cell Ca^2+^ transient

As shown in Fig.2, changing MCU localization can affect the mitochondrial Ca^2+^, but has almost no effect on Ca^2+^ transient and action potential. However, MCU has been reported to be upregulated under disease conditions, such as heart failure (12). Therefore, we performed simulations to further investigate how the maximum MCU activity affected the intracellular Ca^2+^ cycling dynamics. In our simulations, the maximum MCU activity was commanded to increase from control to a higher value after 20 beats (PCL=2 s). Different MCU localizations as in Fig.2 were simulated. Figs.3 A-C shows the time courses of 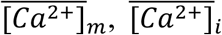, and 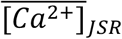 for the three different MCU localizations. The maximum MCU activity was increased 20-fold at t=40 sec, causing mitochondrial Ca^2+^ to gradually increase as expected. However, 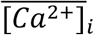 showed a sudden decrease and then gradually increased and eventually reached a value higher than the control for all the three cases. 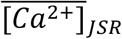 exhibits a similar behavior as the cytosolic Ca^2+^. For comparison, the steady-state peak values (after 50 beats) of Ca^2+^ concentration in mitochondria, bulk cytosol and SR were measured for 1-fold, 10-fold, and 20-fold increases of the maximum MCU activity (Fig.3D). The results show that increasing the MCU activity up to 20-fold resulted in slight increases in the steady-state peak values of 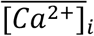 and 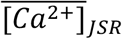, with MCU facing the dyadic space having the most pronounced effect (Fig.3D, cyan). However, as shown in our previous study (12), under heart failure conditions, increasing MCU activity can promote EADs due to the positive feedback loop between Ca^2+^ cycling and action potential.

**Figure 3.**
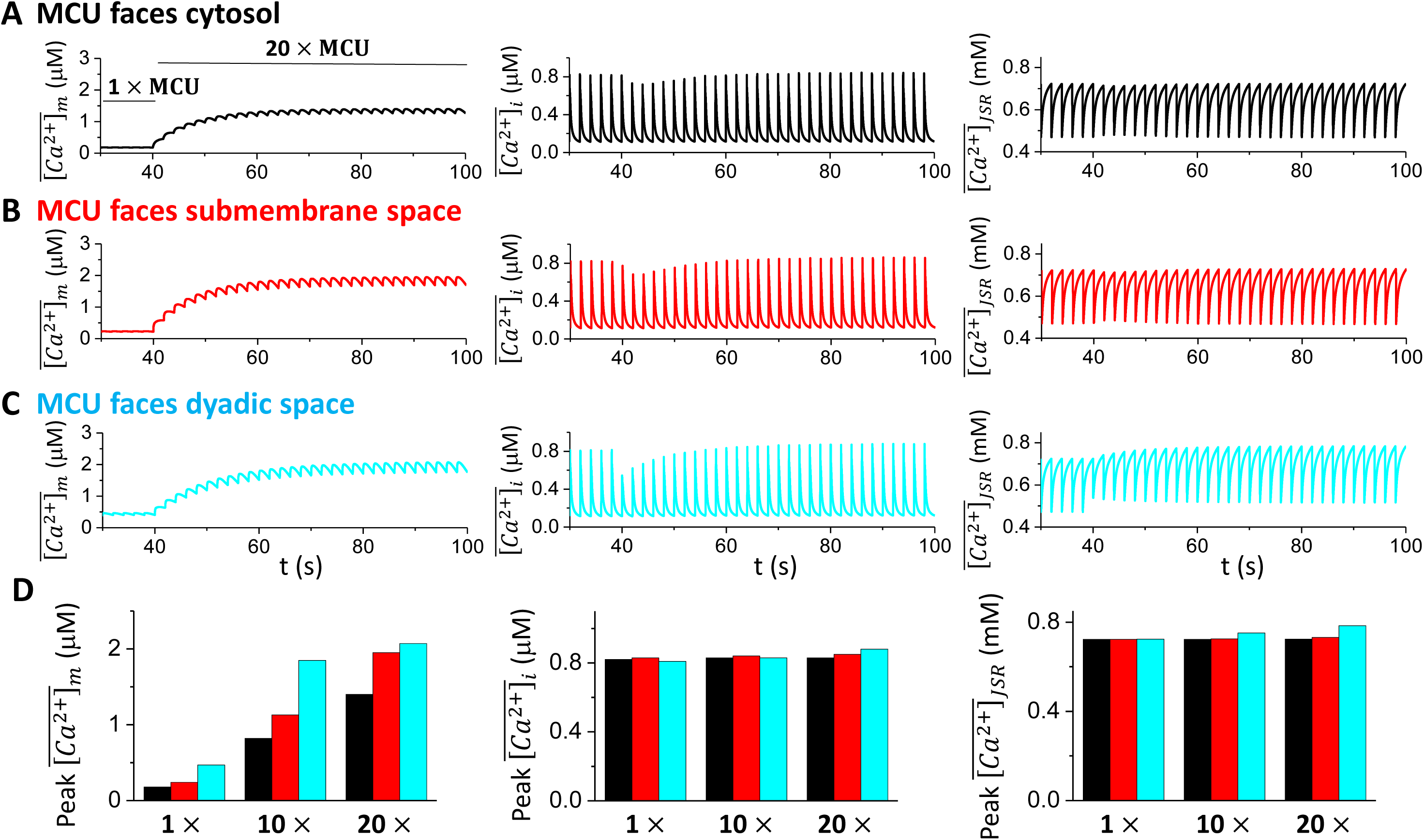
MCU strength on intracellular Ca^2+^ cycling. The maximum MCU activity was increased to a higher level after the cell was paced into the steady state (20 beats). The total number of simulated beats for each case is 50, and the PCL is 2 sec. From left to right, time courses of 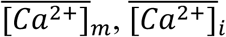, and 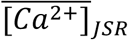. **A**. MCU faces the cytosolic space. **B**. MCU faces the submembrane space. **C**. MCU faces the dyadic space. **D**. Bar plots summarizing the peak values of 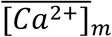 (left), 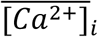 (middle), and 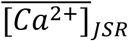 (right) at the 50^th^ beat for different MCU localizations and different maximum MCU activity.

### MPTP and Ca^2+^ cycling behaviors in single mitochondria

We next compared the effects of MPTP opening on Ca^2+^ cycling properties in single mitochondria from our cell model to those reported experimentally by Lu et al. (55). In the simulations, we set V=-80 mV, maintained the diastolic 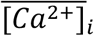 in close to 0.1 μM by tuning the background Ca^2+^ current across the cell membrane, and inhibited NCX_m_ to mimic the experimental condition in Lu et al. (55). We recorded the time when MPTP opens and the opening duration of each event. The histograms of those quantities are shown in Fig.4A along with the experimental results by Lu et al. (55). The transition rates of the Markovian MPTP model are chosen such that the average MPTP open frequency, MPTP open probability, and the open time in the model were 1.4×10^−4^ *mito*^−1^ min^−1^, 0.012%, and 51.49 sec, respectively, which agree approximately with the experimental data (Fig.4B). The time course of [Ca^2+^]_m_ is shown in the right panel of Fig.4C, which agrees with the experimental observations by Lu et al. shown in the left panel of Fig.4C.

**Figure 4.**
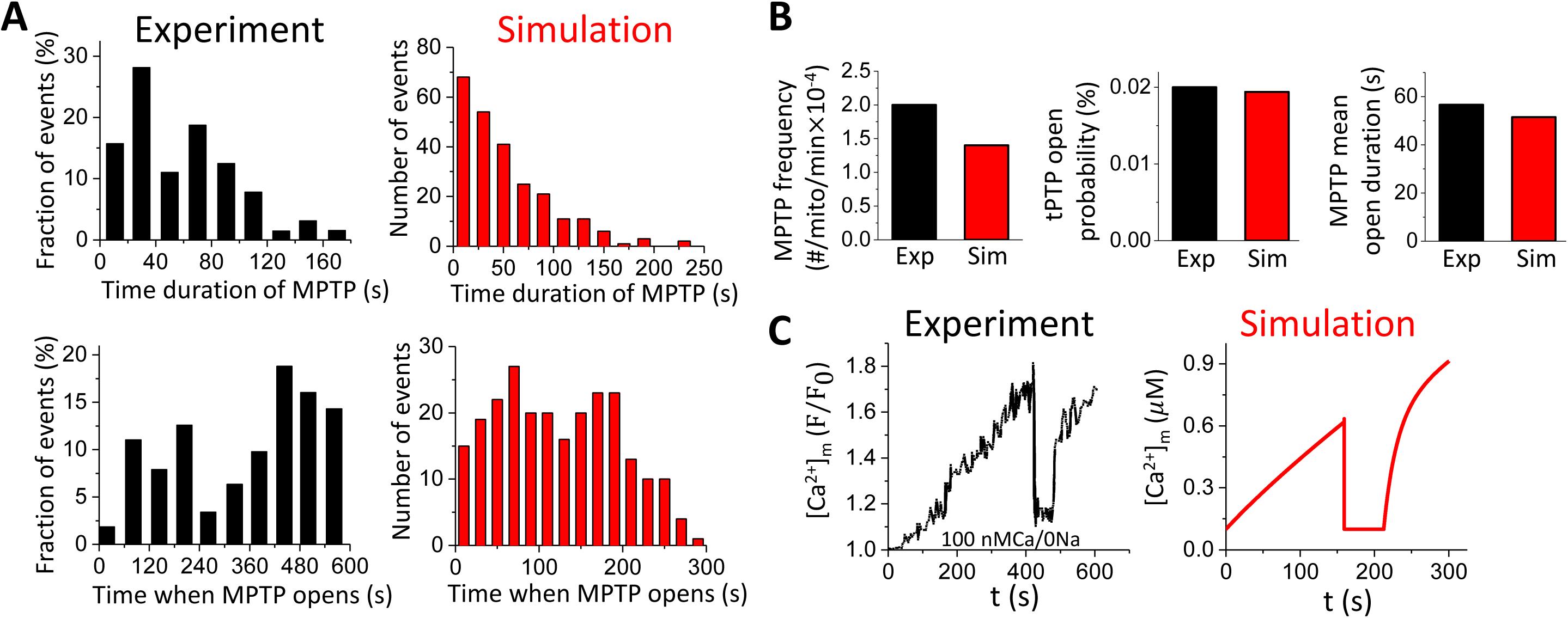
MPTP activity and Ca^2+^ cycling behavior in a single mitochondrion. The membrane potential is clamped at −80 mV. The free cytosolic Ca^2+^ maintains at ∼0.1*μ*M by adjusting the background Ca^2+^ current. The total simulated time is 300 sec for each simulation. **A**. Distributions of MPTP open duration and time when MPTP opens from experiments by Lu et al. (55) and our simulations. **B**. MPTP open frequency, open probability and mean open duration from experiment by Lu et al. (55) and our simulations. **C**. Time course of mitochondrial free Ca^2+^ with MPTP opening and closing from experiment by Lu et al. (55) and our simulation in a single mitochondrion.

### Effects of MPTP opening on Ca^2+^ sparks and the whole cell Ca transient

Experiments have shown that mitochondrial depolarization through MPTP opening exhibits a small effect on Ca^2+^ spark amplitude (∼15%) but dramatically increases spark frequency (∼ 3-fold) (1, 2, 52). Here we followed the protocol by Boyman et al. (52) in which photon stress was used in a region of the cell to generate ROS in the mitochondria, leading to mitochondrial depolarization. In our model, mitochondria were depolarized in a region (Fig.5A) by increasing the close-to-open rate of MPTPs to ensure ∼100% open probability of MPTP within this region. ROS in the depolarized region reached ∼200 *μ*M, and quickly declined outside the region due to short life time of ROS (Fig.5A). Fig.5B shows a line-scan (marked by the red line in Fig. 5A) image of the cytosolic Ca^2+^ and Ca^2+^ profiles from the three zones (I, far away from the depolarized region; I, intermediate region; and III, depolarized region as marked in Fig.5A). The spark frequency was much higher in the depolarized zone, but the spark amplitude remained similar in all the three regions. We compare our results with the experimental results from Boyman et al. (52) in Fig.5 C and D. Fig.5C plots the sample profiles of Ca^2+^ sparks from the three different zones, which are almost identical, in agreement with the experimental recordings by Boyman et al. (52). We then collected Ca^2+^ spark events for 100 seconds, and plotted the Ca^2+^ spark amplitude and frequency from the three different zones (Fig.5D), which shows the amplitude was reduced by ∼15% and the frequency was increased by ∼3-fold for sparks from the depolarized zone (III) compared to those from the far away region (I), also agreeing the experimental results by Boyman et al. (52).

**Figure 5.**
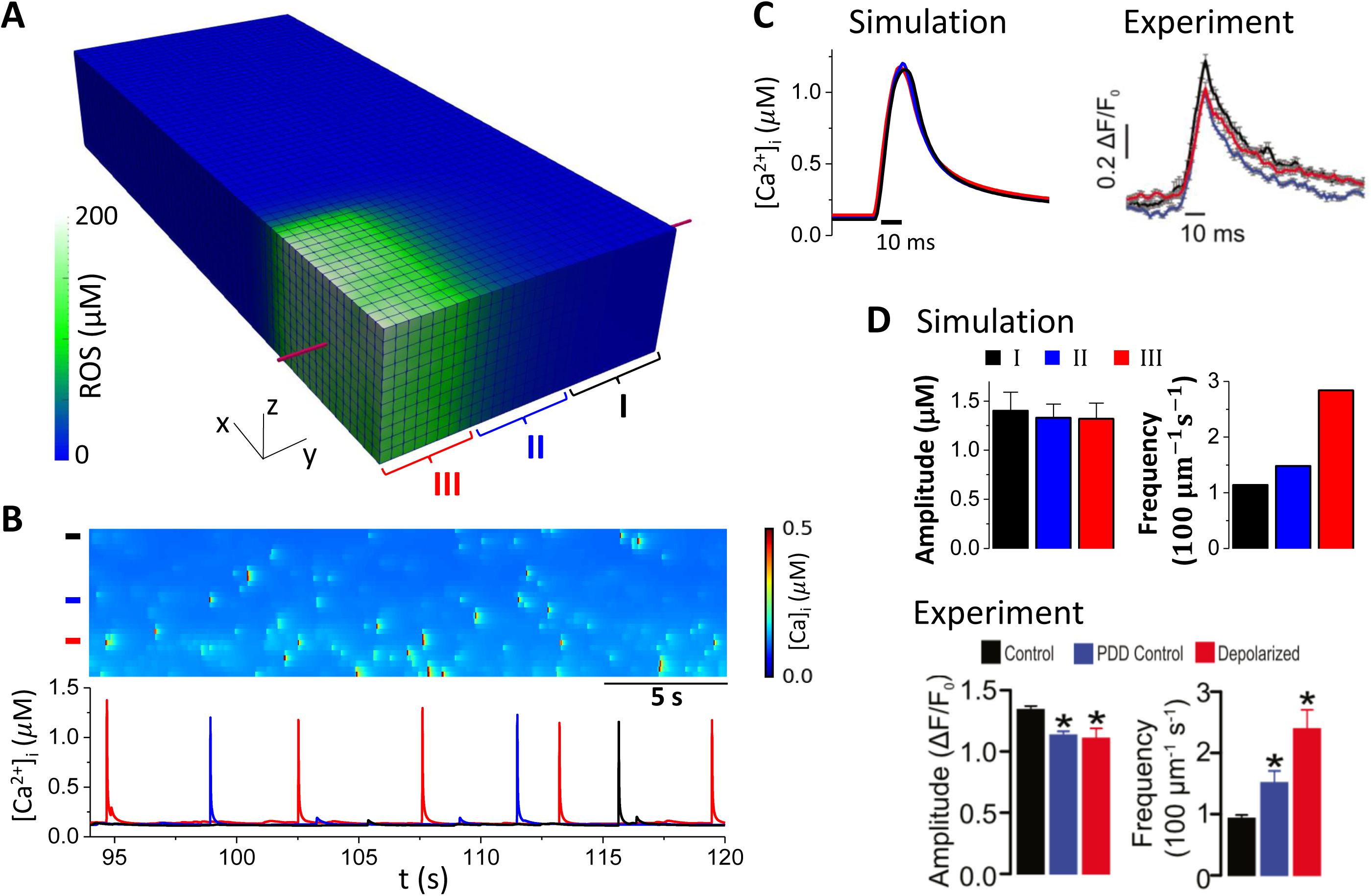
MPTP opening on the properties of Ca^2+^ sparks. The cell remains at rest (V=-80 mV). The MPTPs in the region where n_x_≤16 and n_y_≤6 are depolarized by increasing the MPTP close-to-open rate 2000-fold. The total time simulated is 100 sec. We define three zones similar to those in Boyman et al. (52): I, region that is far from the depolarized region (20<n_y_≤28); II, intermediate region (7<n_y_≤20); III, depolarized region (n_x_≤16, n_y_≤6). **A**. ROS distribution in the cell during mitochondrial depolarization in assigned region. **B**. Top: Line scan image of Ca^2+^ sparks along the red line marked in A. Bottom: Sample Ca^2+^ traces from the three zones as marked by the color bars in the top panel. **C**. Left panel: overlap plot of the Ca^2+^ spark profiles from the three zones. Right panel: experimental measurement of Ca^2+^ spark profiles in different zones by Boyman et al. (52). **D**. Top: Ca^2+^ spark amplitude and frequency measured from our simulations. Bottom: same quantities from experiments by Boyman et al. (52).

The experiments by Boyman et al. (52) showed that Ca^2+^ spark frequency and amplitude remained the same when mitochondria depolarized without ROS elevation, indicating that ROS may play a key role in changing the properties of Ca^2+^ sparks. To test effects of ROS generated by mitochondrial depolarization on Ca^2+^ sparks, we clamped ROS at the control level, i.e., 0.1 *μ*M. The cell was at rest with no pacing. After mitochondrial depolarization, the mitochondrial Ca^2+^ level increased slightly (from 0.11 *μ*M to 0.13 *μ*M), tracking the cytosolic Ca^2+^. However, the whole-cell averaged Ca^2+^ concentration 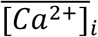 and SR load 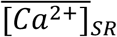 exhibited almost no change after the depolarization (Fig.6A). Both the Ca^2+^ spark amplitude and frequency remained almost the same as control. These results agree with the experimental observation by Boyman et al. (52) that ROS are required to change spark properties. We also carried out simulations with pacing at a PCL of 1 sec (Fig.6B). After mitochondrial depolarization, the whole-cell Ca^2+^ transient 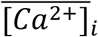 exhibited a transient increase and returned to a new steady state where the peak Ca^2+^ was lower than the control. For comparison, we performed the same simulation but with ROS being clamped to the control level (0.1 *μ*M). Under this condition, the cytosolic Ca^2+^ exhibited similar transient increase after mitochondria were depolarized, but at steady-state, the peak Ca^2+^ was the same as control. These results suggest that the buffered Ca^2+^ inside the mitochondria may transiently affect intracellular Ca^2+^, but it is the ROS that play a key role in altering intracellular Ca^2+^ dynamics during mitochondrial depolarization.

**Figure 6.**
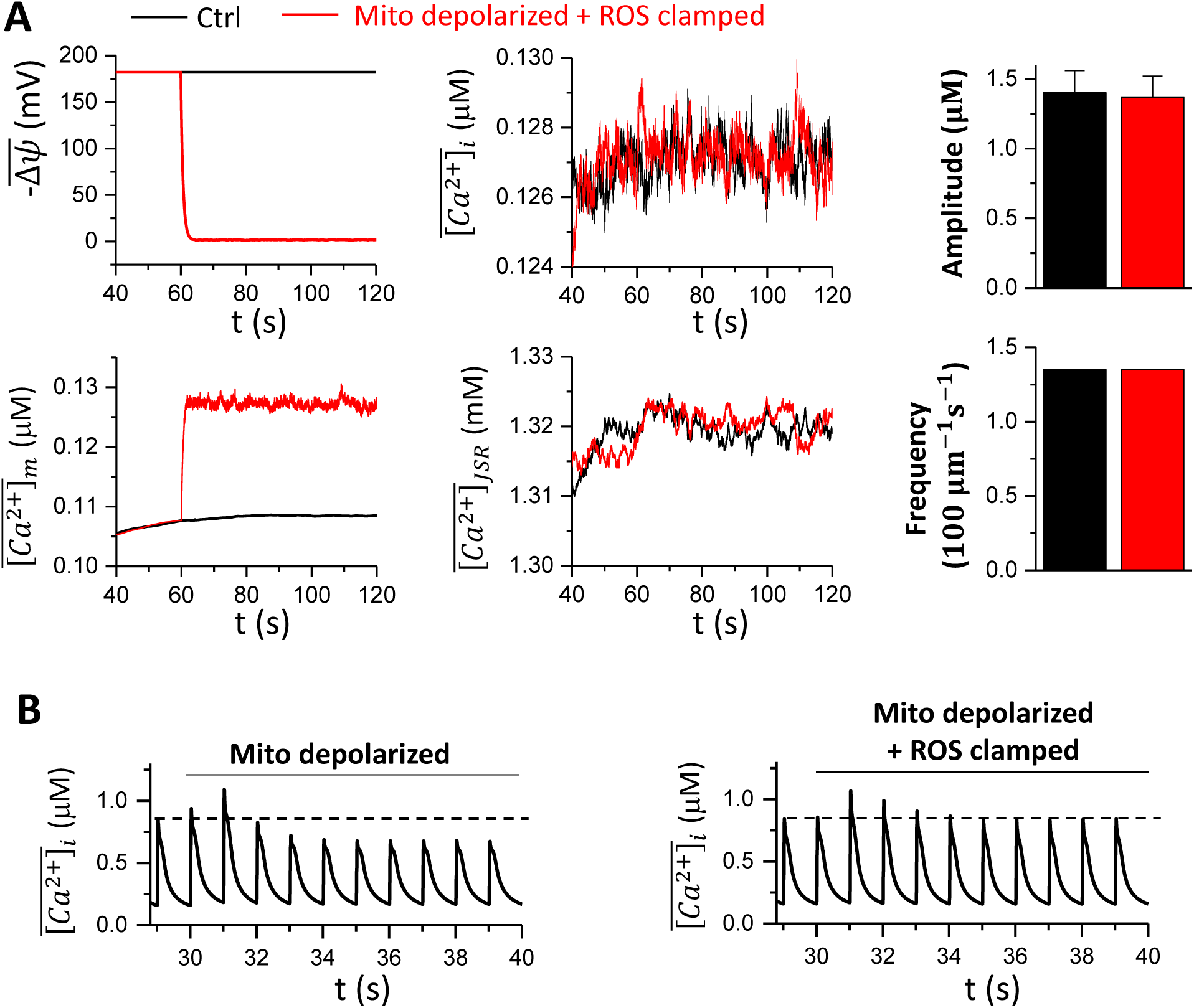
Effects of ROS on Ca^2+^ spark and Ca^2+^ transient behaviors. **A.** Time courses of 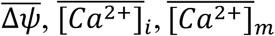, and 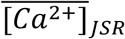 without (black) and with (red) clamped ROS (0.1 µM). The right panels show the corresponding Ca^2+^ spark amplitude and frequency. In the simulations, the cell remained at rest (V=-80 mV). The total time simulated is 120 sec. **B**. The whole cell Ca^2+^ transient during mitochondrial depolarization with MCU sensing the dyadic Ca^2+^ for free running ROS (left) and clamped ROS (right). The cell is paced at PCL 1 sec. The total time simulated is 40 sec.

### Mitochondrial depolarization promotes Ca^2+^ alternans and spontaneous Ca^2+^ waves

Many experimental studies (3-6, 9) have shown that metabolic stress or mitochondrial depolarization can cause Ca^2+^ alternans and spontaneous Ca^2+^ waves. Here we used our model to simulate the effects of mitochondrial depolarization on Ca^2+^ alternans and waves.

To investigate the effect of mitochondrial depolarization on Ca^2+^ alternans, a PCL of 0.5 sec was chosen such that under control conditions, the cell did not exhibit Ca^2+^ alternans. With the cell model paced into the steady state, mitochondria in the cell were depolarized through the commanded opening of MPTP. Immediately after the mitochondrial depolarization, Ca^2+^ alternans developed (Fig.7A, left). However, when the ROS level was clamped to 0.1 µM during the mitochondrial depolarization, Ca^2+^ alternans was not observed (Fig.7A, right).

**Figure 7.**
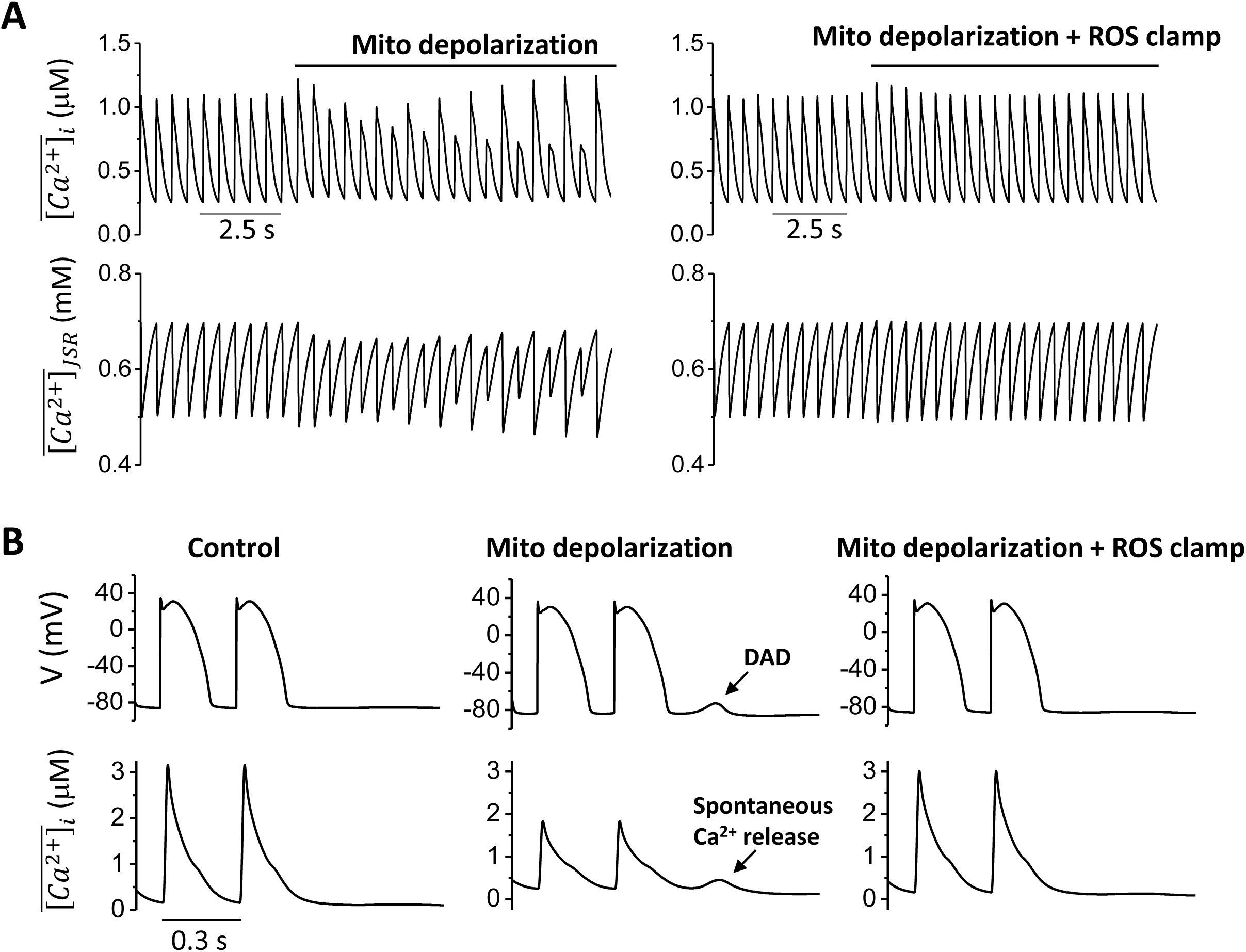
Mitochondrial depolarization promotes Ca^2+^alternans and spontaneous Ca^2+^ waves. **A.** Time courses of 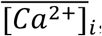, and 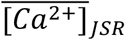 at PCL=0.5 s for free running ROS and clamped ROS (0.1 μM). Mitochondria are depolarized after the cell is paced into the steady state. **B**. Time courses of cell membrane potential (V) and Ca^2+^ transient 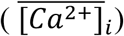 at PCL=0.3 s for control, mitochondrial depolarization with free running ROS, and mitochondrial depolarization with clamped ROS (0.1 μM).

To study the effect of mitochondrial depolarization on spontaneous Ca^2+^ waves and DADs, SERCA pump activity was increased (*v*_*up*_ = 0.8 µM · msV1, and k_i_ = 0.3 *μM*.) and the ROS effect on RyR leakiness was enhanced (Δk_ROS,max_ = 2.). The cell model was paced into the steady state at a PCL of 0.3 sec and then abruptly stopped. When no MPTP opened after pacing, no spontaneous Ca^2+^ release or DADs were observed (Fig.7B, left). However, when MPTP opening occurred such that mitochondria depolarized, spontaneous Ca^2+^ release and DADs did occur after the pacing stopped (Fig.7B, middle), unless ROS levels were clamped to 0.1 µM (Fig.7B, right).

These results show that our model successfully reproduced experimental observations that mitochondrial depolarization promotes Ca^2+^ alternans and spontaneous Ca^2+^ waves. Furthermore, these simulations suggest that intracellular ROS generation in response to mitochondrial depolarization plays a key role in these phenomena.

## Discussion

In this study, we developed a new spatiotemporal ventricular myocyte model that integrates membrane voltage, cytosolic, SR, and mitochondrial Ca^2+^ cycling, as well as ROS and oxidative CaMKII signaling. The model correctly simulates the effects of mitochondrial Ca^2+^ cycling and depolarization on intracellular Ca^2+^ cycling and action potential dynamics observed in experiments. Specifically, we show that under normal conditions, MCU localization can exhibit a large effect on mitochondrial load but has very small effects on intracellular Ca^2+^ cycling and action potential. Altering MCU activity has similar effects. Mitochondrial depolarization via MPTP opening slightly lowers Ca^2+^ spark amplitude but causes a more than 2-fold increase of spark frequency. Mitochondrial depolarization promotes spontaneous Ca^2+^ release causing DADs and can induce intracellular Ca^2+^ alternans. These effects are mediated mainly via ROS signaling since they disappear if ROS levels are clamped.

Computer models of mitochondrial metabolism and its coupling with excitation and contraction have been developed previously (47, 65-72). Some of the models have been used to study the action potential dynamics in single cells (11) and tissue (15). However, most of these models are not spatially extended models, and thus cannot simulate the spatiotemporal aspects of intracellular Ca^2+^ cycling dynamics (9, 10) and mitochondrial depolarization dynamics (26-33). Our model is a spatiotemporal model that can be used to simulate these spatiotemporal dynamics and unravel the dynamical mechanisms arising from coupling of mitochondrial membrane potential and mitochondrial Ca^2+^ cycling, intracellular Ca^2+^ cycling, and action potential of ventricular myocytes. As shown in the present study, our model accurately captures the experimentally observed properties of single Ca^2+^ sparks, Ca^2+^ cycling properties of single mitochondria, and whole Ca^2+^ transient dynamics caused by mitochondria.

Limitations still exist in this model. We ignored detailed glycolytic and metabolic regulation that has been described in detail in other models (47, 65-72), which were replaced with simplified models to describe Δψ, ROS, and ATP. One rationale for this simplification is that the time scale of metabolism is much slower than electrophysiology, so it is not unreasonable to treat them as parameters instead of variables. Detailed models of glycolysis and mitochondrial metabolism have been developed in previous models (32, 50, 51), which can be easily incorporated into this model if desired.

## Author contributions

Z.S. and Z.Q. designed the overall research; Z.S. performed the simulations; L-H.X. and J.N.W. contributed to the overall research design; Z.S., L-H.X., J.N.W., and Z.Q. analyzed data and wrote the article.

## Acknowledgements

This study is supported by NIH grant R01 HL133294.

